# Treating the lungs via the brain: Mechanisms underpinning improvements in breathlessness with pulmonary rehabilitation

**DOI:** 10.1101/117390

**Authors:** Mari Herigstad, Olivia Faull, Anja Hayen, Eleanor Evans, Maxine F. Hardinge, Katja Wiech, Kyle T. S. Pattinson

## Abstract

**Background:** Breathlessness in chronic obstructive pulmonary disease (COPD) is often discordant with airway pathophysiology (“over-perception”). Pulmonary rehabilitation has profound effects upon breathlessness, without influencing lung function. Learned associations can influence brain mechanisms of sensory perception. We therefore hypothesised that improvements in breathlessness with pulmonary rehabilitation may be explained by changing neural representations of learned associations, reducing “over-perception”.

**Methods:** In 31 patients with COPD, we tested how pulmonary rehabilitation altered the relationship between brain activity during learned associations with a word-cue task (using functional magnetic resonance imaging), clinical, and psychological measures of breathlessness.

**Results:** Improvements in breathlessness and breathlessness-anxiety correlated with reductions in word-cue related activity in the insula and anterior cingulate cortex (ACC) (breathlessness), and increased activations in attention regulation and motor networks (breathlessness-anxiety). Greater baseline (pre-rehabilitation) activity in the insula, ACC and prefrontal cortex correlated with the magnitude of improvement in breathlessness and breathlessness anxiety.

**Conclusions:** Pulmonary rehabilitation reduces the influence of learned associations upon neural processes that generate breathlessness. Patients with stronger word-cue related activity at baseline benefitted more from pulmonary rehabilitation. These findings highlight the importance of targeting learned associations within treatments for COPD, demonstrating how neuroimaging may contribute to patient stratification and more successful personalised therapy.

## INTRODUCTION

Breathlessness is an “*all-consuming and life-changing*” (1) experience that is subjective, intensely personal (2) and is associated with profound fear. It is the primary symptom of chronic obstructive pulmonary disease (COPD). Importantly, the severity of reported breathlessness is frequently discordant with measures of airway pathophysiology (3). The terms “over-perception” and “under-perception” are often used to describe discordant breathlessness.

One of the most effective treatments for breathlessness in stable COPD is pulmonary rehabilitation (4). It benefits both personal wellbeing and exercise capacity, but has no measurable effect on lung function (5). However, the clinical response to pulmonary rehabilitation is variable. Although 50-60% of participants improve, approximately 40% of people who complete the treatment derive no measurable benefit (6). The cause for this variation in treatment response remains unknown. A better understanding of the mechanisms underlying breathlessness perception would help us comprehend the discordance with measures of airway pathophysiology, and why the response to pulmonary rehabilitation is so variable. Such understandings would help to personalise therapy for COPD.

Recently there has been a step-change in the neuroscientific understanding of sensory perception (2, 7), emphasising the importance of top-down perceptual processing. This has important implications for the understanding of breathlessness. The traditional bottom-up model of stimulus perception theorises that ascending sensory information (e.g. vagal afferents from the airways) is modulated by brain processes such as learning, expectation, and affect, ultimately generating the conscious perception of the sensation (e.g. breathlessness) (8). Recent models suggest that sensory perception includes a more prominent top-down process (7, 9), in which experienced sensations are derived “from the brain’s predictions about the causes of events in the body, with incoming sensory inputs keeping predictions in check (9).” Therefore, instead of being a stimulus-response organ that passively waits for ascending inputs, the brain forms predictions constructed from previous experiences (otherwise known as priors). These priors are then updated or corrected by neural observations of incoming sensory information (7, 9). Negative affect, attention and interoceptive ability may act as moderators within this system, adjusting either the priors or weighting (gain) of incoming sensory information to increase symptom discordance (or “over/under perception” of symptoms).

The neurobiological basis for generating priors is thought to reside in a stimulus valuation network (9) that comprises the anterior insula, anterior cingulate cortex (ACC), orbitofrontal cortex (OFC) and ventromedial prefrontal cortex (9). The stimulus valuation network generates information and predictions about bodily state and emotion. Incoming respiratory sensory information is then fed into this network from the vagus nerve, via the periaqueductal gray (PAG) (10, 11) and posterior insula.

In the context of COPD, repeated episodes of breathlessness reinforce learned associations between context-relevant cues (e.g. stairs, steep hills) and being breathlessness. These learned associations then influence the brain’s set of priors. Strong priors may begin to dominate perception of sensations, accompanied by reduced gain in sensory processing regions. Together, these processes facilitate decoupling of symptoms from objective physiology, promoting “over-perception” of breathlessness. This decoupling is further supported by negative affect such as depression and anxiety (12). Resulting worsening breathlessness can then drive a downward spiral of activity avoidance and physical deconditioning that ultimately leads to further worsening breathlessness.

Pulmonary rehabilitation interrupts this downward spiral of decline. Participants learn to cope better with their breathlessness through repeated exposure to exercise-induced breathlessness in a ‘safe’ healthcare setting. We therefore hypothesised that an important effect of pulmonary rehabilitation would be to reduce learned negative associations with breathlessness-related cues (i.e. a reduction in “over-perception”). This would be reflected in reduced activation in the stimulus valuation network. Extending this hypothesis, we predicted that those subjects with the strongest learned associations might have the most to gain from pulmonary rehabilitation. We therefore assessed conditioned breathlessness and associated brain activity in a group of patients with COPD before and after a course of pulmonary rehabilitation. With functional magnetic resonance imaging (FMRI) we measured the brain’s response to a validated set of breathlessness-related word cues (13) in combination with detailed clinical and psychological characterisation.

## METHODS

A brief overview of the study methods is provided here. Detailed methods are provided at the end of this manuscript. Comparison of pre-rehabilitation FMRI findings with healthy controls and a detailed description of the breathlessness-cue task have been published elsewhere (13, 14).

### Participants

Thirty-one people with COPD (21 male, age 68 years ± 9 (SD)) were studied immediately preceding and following a 6-week course of outpatient pulmonary rehabilitation, delivered by an experienced community pulmonary rehabilitation team. The full course ran for 6 weeks, with two sessions per week including an hour of exercises and an hour of education, as part of a standard pulmonary rehabilitation programme. Patients had been referred to pulmonary rehabilitation as part of their standard management. All participants gave written, informed consent, and the Oxfordshire Research Ethics Committee A approved the study.

### Behavioural and physiological measurements and analysis

#### Self-report questionnaires

The following self-report questionnaires were completed and scored according to their respective manuals: Center for Epidemiologic Studies Depression Scale (CES-D) (15), State-Trait Anxiety Inventory (16), Fatigue Severity Scale (17), St George’s Hospital Respiratory Questionnaire (SGRQ) (18), Medical Research Council (MRC) breathlessness scale (19), Dyspnoea-12 (D-12) questionnaire (20), Catastrophic Thinking Scale in Asthma (modified by substituting the word “breathlessness” for the word “asthma”) (21), Pain Awareness and Vigilance Scale, (modified by substituting the word “breathlessness” for the word “pain”) (22), Behavioral Inhibition System/Behavioral Activation System scale (23).

#### Physiology

Spirometry and an incremental shuttle-walk test were undertaken according to standard practices (24).

#### Analysis

Full correlation matrices were calculated for all behavioural and physiological measures at baseline and for the change following pulmonary rehabilitation, using MATLAB (R2013a, Mathworks, Natick, MA). Partial correlation matrices were calculated on correlated variables, defined as *p* < 0.05 (uncorrected). D12 score was taken as the clinical measure of breathlessness, and was compared to VAS breathlessness scores both prior to and across pulmonary rehabilitation. To explore whether any of our measured behavioural variables were mediating these VAS-D12 relationships, we conducted a mediation analysis using the CANlab mediation toolbox (25).

### Brain imaging and analysis

Magnetic resonance imaging (MRI) was performed with a Siemens 3 Tesla TIM-Trio scanner, using a 12-channel head coil. A structural T1-weighted scan (voxel size 1×1×1 mm), functional (FMRI) T2*-weighted scan (voxel size 3×3×3 mm) and fieldmaps were collected. During the FMRI scans, participants were shown a randomised set of breathlessness-related word cues (13) and asked to rate each cue according to breathlessness and breathlessness-anxiety on a visual analogue scale (VAS), with the question “How breathless would this make you feel?” (wB) and the “How anxious would this make you feel?” (wA). A control condition of matched random letter string presentations were interspersed with breathlessness cues.

#### FMRI Data Analysis

FMRI data processing was carried out within FSL (Oxford Centre for Functional Magnetic Resonance Imaging of the Brain Software Library), using FEAT (FMRI Expert Analysis Tool, Version 5.98). The cluster Z threshold was 2.3 and the corrected cluster significance threshold *p* = 0.05 (26) corrected for multiple comparisons across the whole-brain.

The first-level (individual subject) analysis in FEAT incorporated a general linear model, with explanatory variables (EVs) included for word presentation, and two (de-meaned) EVs modelling the trial-by-trial variability in VAS responses to the breathlessness (wB) and breathlessness-anxiety (wA) cues presented during scanning. A middle-level analysis was then conducted to calculate a difference in brain activity from pre- to post-rehabilitation for each subject.

The group-level analysis aimed to interrogate which brain regions could account for between-subject variability in the VAS responses over the course of pulmonary rehabilitation. To do this, we regressed each individual’s change in wB and wA VAS scores against the change in mean word-cue EV from the middle-level analysis. We also conducted a secondary analysis, where we regressed the change in VAS responses on pre-rehabilitation word-cue brain activity, to probe baseline factors associated with improved breathlessness following pulmonary rehabilitation. Mediation analyses were also conducted on the relationship between VAS score and correlated brain regions, with the behavioural questionnaire scores as potential mediators, using the CANlab mediation toolbox (25).

## RESULTS

### Participants

Demographic, physiological and questionnaire data, as well as averaged breathlessness VAS ratings are presented in Tables 1 and 2. Improvement above the minimally important clinical difference (MCID) in quality of life (as measured by SGRQ (27)) occurred in 16 subjects (52%; MCID = 4 points), functional exercise capacity (as measured by incremental shuttle walk test (28)) in 19 subjects (61%; MCID = -48m), and clinical measures of breathlessness (as measured by D12 (29)) in 14 subjects (45%; MCID = 3 points). Pulmonary rehabilitation led to a group mean improvement in wA VAS responses (pre value 40.5(±SD 19.4) vs. post value 28.8(21.1); *p* = 0.002). Improvements in wB VAS response did not reach statistical significance (pre value 52.6(SD 14.2) vs. post value 49.1(19.9); *p* = 0.086) (Figure 1).

**Table 1:** Participant details and physiological data

|  | Pre<br>rehabilitation | Post<br>rehabilitation | p value |
| --- | --- | --- | --- |
| BMI (kg/m<sup>2</sup>) | 27.8 (5.8) |  |  |
| Smoking (pack years) | 40(35) |  |  |
| Resting Borg score (1-10) | 0.7 (1.0) | 0.4 (0.5) | 0.187 |
| Resting SaO<sub>2</sub> (%) | 94.6 (2.8) | 94.2 (3.1) | 0.313 |
| Resting HR (bpm) | 81.4 (13.0) | 81.4 (14.2) | 1.0 |
| Post-exercise Borg score (1-10) | 4.7 (1.3) | 4.5 (1.7) | 0.187 |
| Post-exercise SaO<sub>2</sub> (%) | 89.4 (6.1) | 89.2 (6.7) | 0.755 |
| Post exercise HR (bpm) | 112.2 (23.0) | 120.7 (24.5) | 0.002 |
| FEV<sub>1</sub> (% predicted) | 73.5 (19.2) | 75.7 (29.9) | 0.695 |
| FEV<sub>1</sub>/FVC % | 58.0 (15.9) | 59.6 (23.2) | 0.723 |

*Demographic data (mean results and standard deviation). MRC score (Medical Research Council breathlessness score, median (interquartile range)), BMI (body mass index), SaO<sub>2</sub> (blood oxygen saturation), MSWT (modified shuttle walking test), FEV<sub>1</sub> (forced expiratory volume in 1 second).*

**Table 2:** Questionnaire and VAS scores

| **Questionnaire** | **Visit 1**<br>**(mean (SD))** | **Visit 2**<br>**(mean (SD))** | **p value** |
| --- | --- | --- | --- |
| MRC score (1-5) | 2.9 (0.8) | 2.7 (0.9) | 0.231 |
| Dyspnoea-12 | 12.0 (9.2) | 7.9 (6.4) | 0.009 |
| SGRQ | 49.9 (17.4) | 42.7 (14.2) | 0.001 |
| Catastrophic<br>Thinking Scale <sup>a</sup> | 12.1 (10.3) | 7.2 (5.4) | 0.022 |
| Awareness and<br>Vigilance Scale <sup>b</sup> | 40.5 (13.0) | 39.0 (14.2) | 0.514 |
| Fatigue Severity<br>Scale | 41.4 (11.3) | 32.6 (11.6) | 0.001 |
| CES-D | 13.3 (8.9) | 11.8 (7.3) | 0.375 |
| State anxiety | 35.1 (9.1) | 32.3 (9.4) | 0.150 |
| Trait anxiety | 35.8 (9.1) | 31.9 (8.8) | 0.010 |
| BIS / BAS | 54.0 (7.4) | 53.9 (7.2) | 0.906 |
| ISWT (m) | 342 (199) | 426 (212) | 0.000001 |

Behavioural measurements before and after pulmonary rehabilitation (n=31, mean results and standard deviation). SGRQ (St George's Respiratory Questionnaire), CES-D (Center for Epidemiologic Studies Depression Scale). ISWT incremental shuttle walk test, wB VAS scale "*How breathless would this make you feel*", wA VAS scale "*How anxious would this make you feel?*"

**Figure 1.**
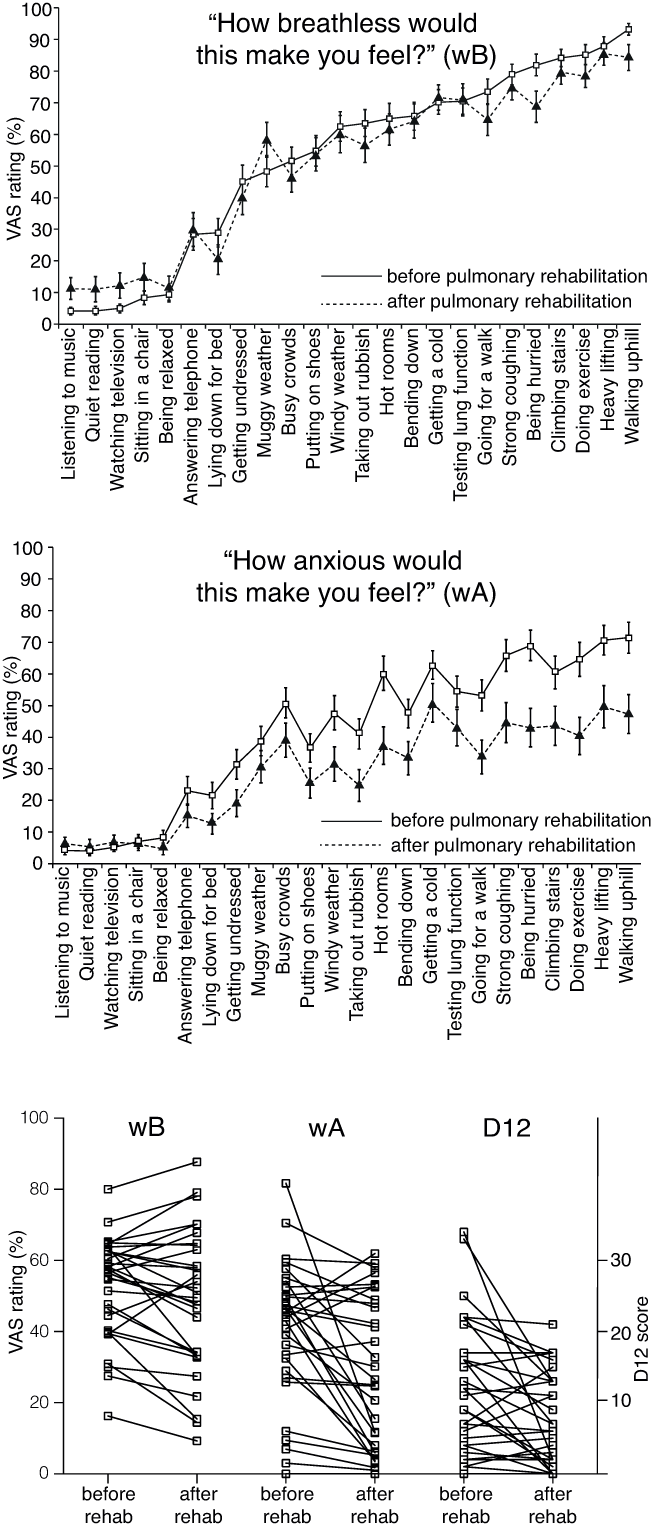
Demonstration of changes in VAS scales relating to “*How breathless would this make you feel?*” (wB; upper figure) and “How anxious would this make you feel” (wA; middle figure), and individual changes in wB, wA and Dyspnea-12 (D12) score over the course of pulmonary rehabilitation (lower figure).

### Variability of response to pulmonary rehabilitation (Figure 2)

#### “How breathless would this make you feel?” (wB)

The greater the reduction in wB over the course of pulmonary rehabilitation, the greater was the reduction in BOLD activity in the left anterior insula, the left posterior insula, the left supramarginal gyrus and the anterior cingulate cortex (i.e. a positive correlation).

**Figure 2.**
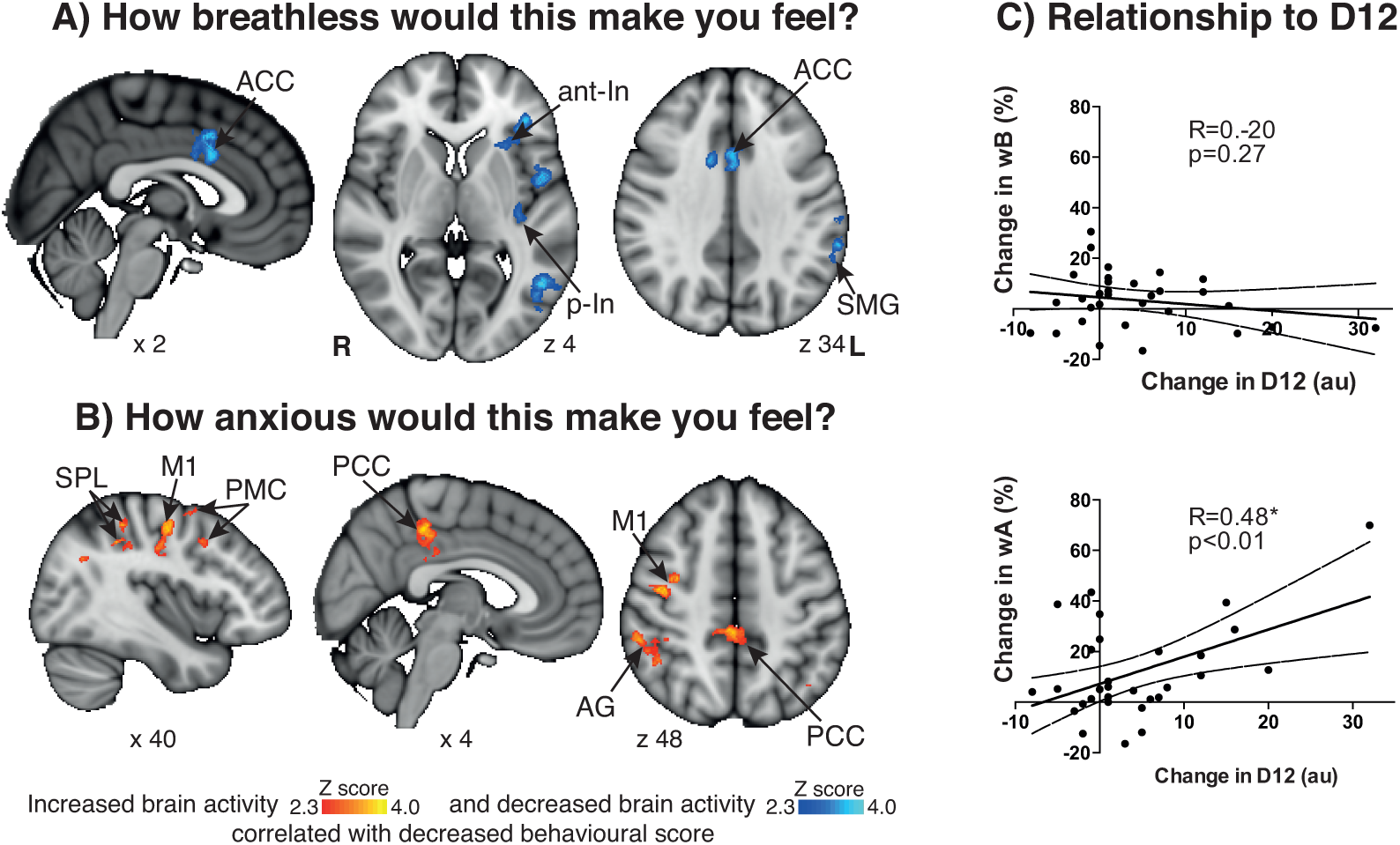
Change in BOLD activity that correlates with rehabilitation-induced changes in behavioural scores of breathlessness response to “*How breathless would this make you feel?*” (wB) (blue-light blue) and “How anxious would this make you feel”(wA) (red-yellow). The images consist of a colour-rendered statistical map superimposed on a standard (MNI 1×1×1 mm) brain, and significant regions are displayed with a threshold Z > 2.3, with a cluster probability threshold of *p* < 0.05 (corrected for multiple comparisons). Abbreviations: ACC, anterior cingulate cortex; PCC, posterior cingulate cortex; ant-In, anterior insula; p-In, posterior insula; SMG, supramarginal gyrus, AG, angular gyrus; PMC, premotor cortex; M1, primary motor cortex; SPL, superior parietal lobe; L, left; R, right. On the right of the image is displayed the linear regression (and 95% confidence intervals) between wB and wA with change in D12 score.

#### “How anxious would this make you feel?” (wA)

The greater the reduction in wA over the course of pulmonary rehabilitation, the greater was the increase in BOLD activity in the posterior cingulate cortex, the angular gyrus, the primary motor cortex and the supramarginal gyrus (i.e. a negative correlation).

#### Behavioural changes with pulmonary rehabilitation

The D12 score was taken to represent a clinically relevant measure of breathlessness. A full correlation matrix across all behavioural scores (Figure 3) showed that change in D12 with pulmonary rehabilitation significantly correlated with the change in wA but not wB from the breathlessness task. Change in D12 also significantly correlated with the St George’s Respiratory Questionnaire, and breathlessness-related catastrophising and vigilance, as well as depression, state and trait anxiety (Figure 3). Using a partial correlation of these variables, only breathlessness-related catastrophising and depression remained independently correlated with D12 (Figure 3).

**Figure 3.**
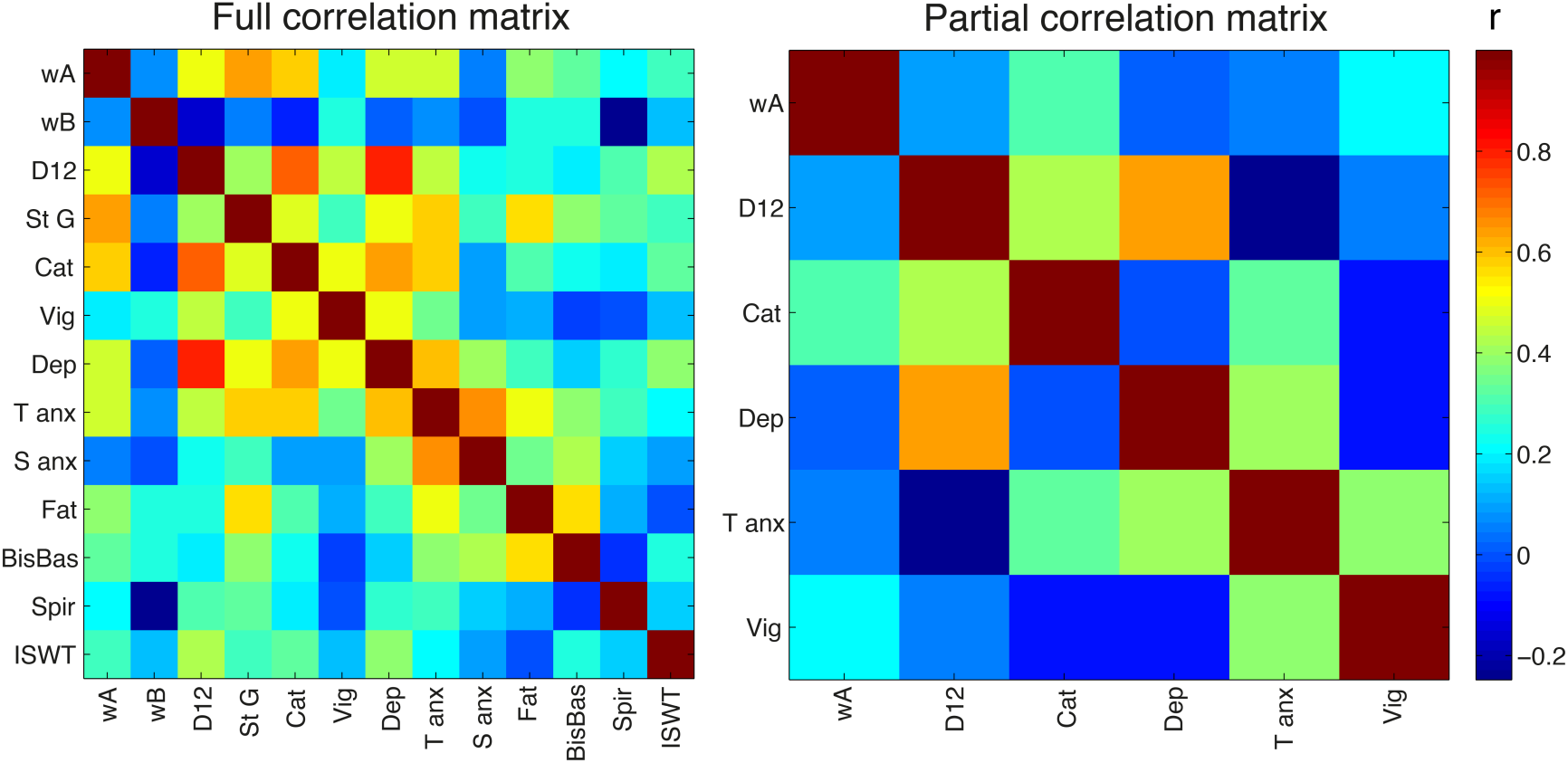
Left: Full correlation matrix of all the measured behavioural variables. Right: Partial correlation matrix of those variables that significantly correlated with the clinical measure of breathlessness (D12, defined at *p* < 0.05, uncorrected). The St George’s Respiratory Questionnaire and incremental shuttle walk test were excluded from the partial correlation despite significant correlation, as they are summary measures combining various psychological constructs. Abbreviations: wA, visual analogue scale (VAS) “How anxious would this make you feel?”; wB, VAS “How breathless would this make you feel?”; D12, Dyspnoea-12 questionnaire; St G, St Georges Respiratory Questionnaire; Cat, Breathlessness catastrophising score; Vig, Breathlessness Vigilance and Awareness score; Dep, Depression; T Anx, Trait Anxiety; S Anx, State Anxiety; Fat, Fatigue; BisBas, Behavioural inhibition behavioural activation scale; Spir, Spirometry (FEV1/FVC); ISWT, incremental shuttle walk test.

#### Mediation analysis

Using a mediation analysis (25) to investigate the relationship between our task-associated change in wA and its relationship to change in D12, we show that this was significantly mediated by change in depression, and the residuals (once depression was accounted for) were non-significant (Figure 4).

**Figure 4.**
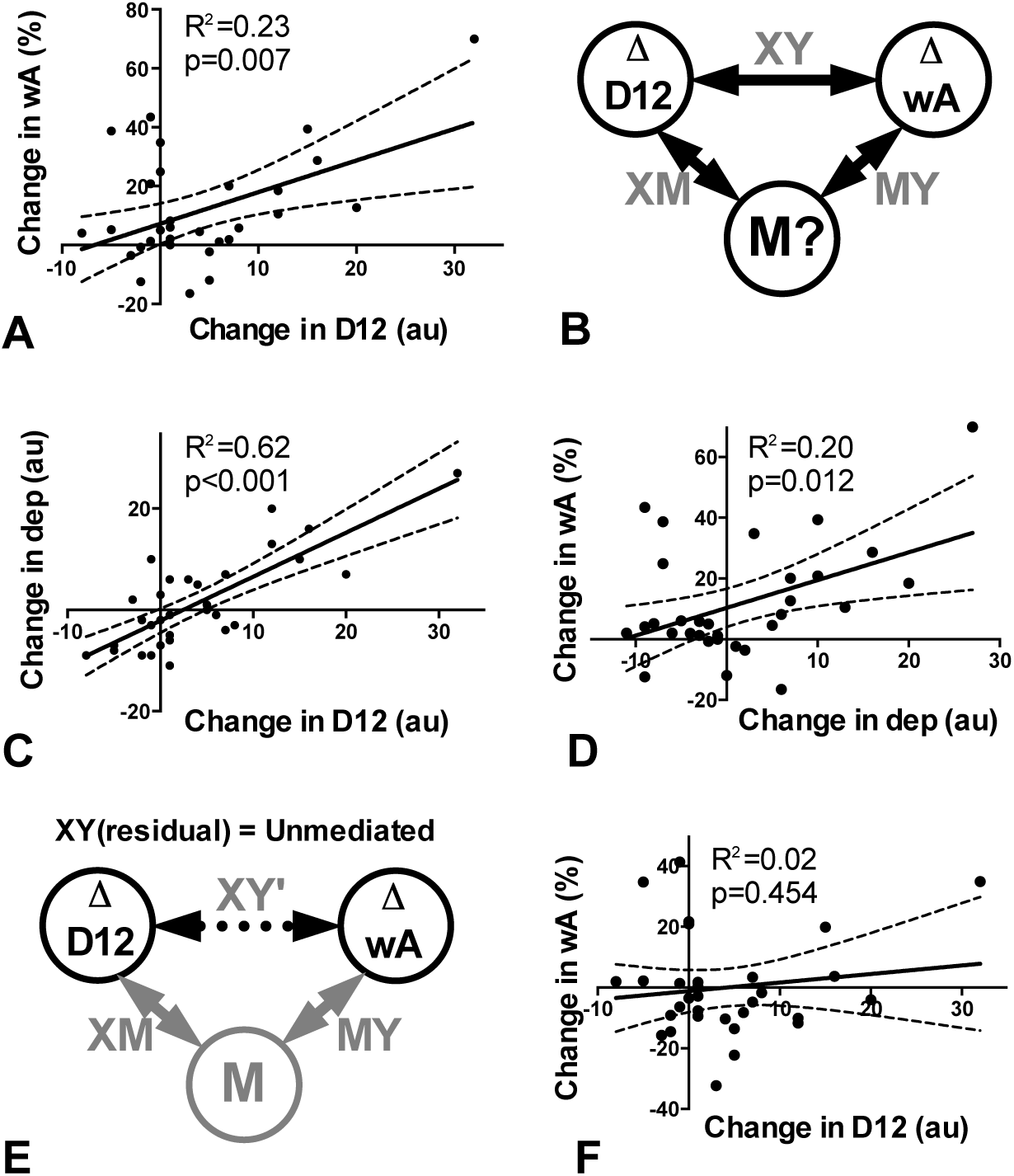
Mediation analysis. Relationship between change in word-cue breathlessness anxiety (wA) and change in D12 questionnaire across the course of pulmonary rehabilitation, mediated by change in depression score. A) Correlation between change in VAS score for wA and change in D12 score. B) Depiction of a mediation, where the relationship between change in wA and D12 (XY) can be mediated by M, which correlates with both X (D12; XM) and Y (wA; XY). C) XM relationship: X variable (D12) predicts mediator variable (depression). D) MY relationship: Mediator variable (depression) predicts Y variable (wA). E) Depiction of residual relationship (XY’) once the effect of the mediator variable has been removed. F) Residual relationship between change in wA and change in D12 when change in depression has been regressed out. For full explanation of mediation please see the online supplement.

### Baseline factors associated with improved breathlessness following pulmonary rehabilitation (Figure 5)

#### “How breathless would this make you feel?” (wB)

We observed a positive correlation between change in wB and pre-rehabilitation activity in the anterior insula (bilateral), orbitofrontal cortex (bilateral), precuneus, lateral occipital cortex and primary motor cortex (bilateral), i.e. the stronger the activation at baseline, the greater the reduction in wB score.

#### “How anxious would this make you feel?” (wA)

We observed a positive correlation between change in wA and pre-rehabilitation activity in the anterior cingulate cortex and ventromedial prefrontal cortex, i.e. the stronger the activation at baseline, the greater the reduction in wA score.

**Figure 5.**
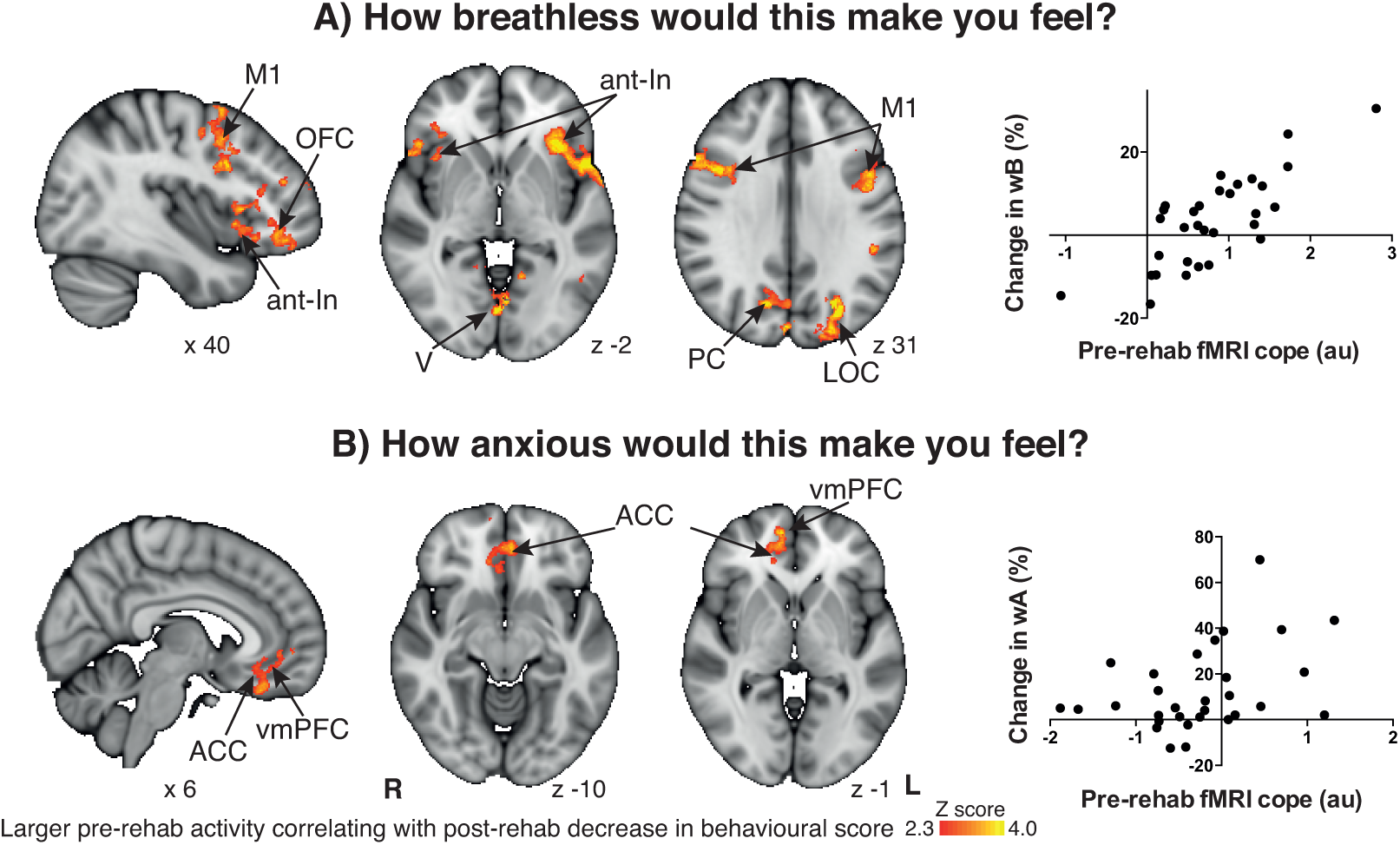
Pre-rehabilitation BOLD activity that correlates with rehabilitation-induced changes in behavioural scores of breathlessness response to “*How breathless would this make you feel?*” (wB) (upper figure) and “How anxious would this make you feel” (wA) (lower figure). The images consist of a colour-rendered statistical map superimposed on a standard (MNI 1x1x1 mm) brain, and significant regions are displayed with a threshold Z > 2.3, with a cluster probability threshold of *p* < 0.05 (corrected for multiple comparisons). Abbreviations: vmPFC, ventromedial prefrontal cortex; ACC, anterior cingulate cortex; OFC, orbitofrontal cortex; PC, precuneus cortex; ant-In, anterior insula; V, visual cortex; LOC, lateral occipital cortex; M1, primary motor cortex; L, left; R, right. The graphs represent linear regression between the averaged COPE value in the significant regions, and the change in wB and wA scores respectively.

## DISCUSSION

### Key findings

Overall, pulmonary rehabilitation delivered expected improvements in breathlessness, exercise capacity, and quality of life in line with previously published research (4, 6, 30, 31). While pulmonary rehabilitation induced a significant overall improvement in many psychological and breathlessness perception measures (Table 2), a range in treatment response was found between individuals. According to the magnitude of treatment response in VAS breathlessness and breathlessness-anxiety scores, pulmonary rehabilitation led to reduced activity in the brain’s stimulus valuation network and increased activity in attention regulating networks. At baseline (i.e. before pulmonary rehabilitation), activity in the stimulus valuation network correlated with change in wB and wA scores following pulmonary rehabilitation, (i.e. the greater the brain’s response to word cues before rehabilitation, the greater the reduction in wB and wA scores after rehabilitation).

### Effect of pulmonary rehabilitation on brain and behaviour

Group improvements were observed in a range of affective domains, including breathlessness-specific measures and generalised anxiety and fatigue (Table 2). These changes occurred independently of any changes in spirometry (5). However, not all participants showed clinically important improvements, emphasizing the need to understand the brain processes that are targeted by pulmonary rehabilitation.

In chronic diseases such as COPD, prior experiences of symptoms can vastly alter outcomes and symptom perception (32). Furthermore, the learned associations between environmental cues and symptoms can influence this symptom expectation and subsequent perception. We therefore used a word-cue task to probe these associations. The improvement in word-cue breathlessness-anxiety (wA) reflected the changes in the clinical measure of breathlessness (D12) across the course of rehabilitation (Figure 3). This suggests that pulmonary rehabilitation may have a greater effect upon the affective interpretation of breathlessness rather than its objective experience as improvements in wB did not reach statistical significance at a group level.

We then examined the changes in brain activity associated with word-cue presentation across pulmonary rehabilitation. When FMRI activity was compared to each individual’s reductions in ratings of breathlessness (wB), correlated reductions in brain activity were observed in the anterior insular cortex, anterior cingulate cortex, prefrontal cortex and posterior insular cortex. Together the anterior insula, ACC, and prefrontal cortex form key components of the stimulus valuation network, which is responsible for the conscious awareness of the internal state of the body, or interoception (9). This observed reduction in activity in this stimulus valuation network may represent re-evaluated associations, towards more appropriate evaluation of breathlessness-cues. In other words, these findings may represent a dampening in activity in brain areas responsible for expectations of breathlessness.

The posterior insula plays a role in sensory-discriminative processing, with connectivity both to somatosensory cortices and to the anterior insula to integrate sensory inputs with the stimulus valuation network (9). The reduced activity seen in this structure relating to reduced wB following pulmonary rehabilitation suggests that the reappraisal of learned associations may also influence lower order-sensory processing.

When FMRI activity was negatively correlated with individual reductions in ratings of breathlessness-anxiety (wA) in the angular gyrus, supramarginal gyrus and posterior cingulate cortex. These changes in brain activity could represent a shift in expectancy towards that seen in healthy controls (14), (33). The angular gyrus and supramarginal gyrus form a multimodal complex that integrates somatosensory inputs to the brain and is associated with attention processing. The posterior cingulate cortex is thought to mediate interactions between emotions and memory, controlling attentional focus (34). All these functions are impaired by anxiety (35, 36). Additionally, the increase in activity in the granular primary and pre-motor cortices may reflect a shift towards more objective perceptual processing less dominated by learned associations (9, 33, 37).

### Baseline FMRI correlates with changes in VAS scores following pulmonary rehabilitation

In our second analysis, we demonstrated that greater improvements in VAS responses with pulmonary rehabilitation were associated with greater BOLD activity in the anterior insula, the ventromedial prefrontal cortex and anterior cingulate cortex (components of the stimulus valuation network) at baseline (Figure 3). These findings imply that people with stronger learned associations between contextual cues and their breathlessness are more likely to benefit from pulmonary rehabilitation. It is possible that the repeated episodes of breathlessness experienced in a “safe” setting during pulmonary rehabilitation help to re-shape associations, which are then reinforced by reappraisal of these situations between the rehabilitation sessions. These findings and interpretations are supported by a recent behavioural study (38), which showed that higher breathlessness-fear before pulmonary rehabilitation was associated with greater reductions in breathlessness during exercise following pulmonary rehabilitation.

### Driving forces behind changes in behaviour

We also interrogated the relationship between changes in the VAS scales and associated brain activity with clinical and behavioural measures of breathlessness. The full correlation matrix (Figure 3) demonstrates that many of the behavioural measures are at least partially explained by each other. A notable exception is spirometry, which is poorly explained by all of the other measures. This distinction between lung function and the various measures of breathlessness perception adds to the weight of evidence that the impact of COPD is poorly reflected by spirometry (3).

When investigating the VAS scores associated with the breathlessness expectancy task, the strongest correlation with wA was the D12 questionnaire; a well validated clinical measure of breathlessness (20). In addition, strong correlations exist between wA and breathlessness catastrophising and depression. Conversely, the wB measure did not correlate significantly with any of the measures, although a trend was observed with change in spirometry (*p* = 0.10). Further analysis (25) revealed that the relationship between wA and the D12 scale appears to be mediated by changes in depression, and when this component is removed there is no remaining relationship between D12 and wA (Figure 4F). Although the direction of the relationship between anxiety, depression and breathlessness remains unknown, many consider it to be bidirectional (12). Some even speculate that aberrant predictions mediated by the stimulus valuation network may drive depression, anxiety and fatigue (9).

Thus, based upon a Bayesian model of sensory perception, we speculate that pulmonary rehabilitation exerts its benefit by two broad mechanisms in the brain: First by updating of the brain’s set of breathlessness-related priors (through associative learning), and second by acting upon measures of negative affect that act as moderators of priors and influence gain of sensory information processing. Improvements in chest wall or quadriceps muscle function with fitness (4) would additionally adjust afferent inputs. These factors are likely to work together to contribute to the well-documented improvements in wellbeing with pulmonary rehabilitation (4, 31, 38).

### Clinical relevance

This study has examined brain mechanisms underlying the variable response to pulmonary rehabilitation. We have shown that improvements in breathlessness over the course of pulmonary rehabilitation are associated with reductions in activity in the stimulus valuation network, and increased activity in attention processing and motor control areas of the brain. We have also demonstrated that improvements responses to the breathlessness-anxiety word cue (wA) correspond with a clinically meaningful measure of breathlessness, the D12 questionnaire. Therefore, specifically targeting associative learning might be a way to further improve efficacy of pulmonary rehabilitation. This might arise from combining pulmonary rehabilitation with drugs that target relevant neurotransmitter systems (39). As many patients decline the invitation to pulmonary rehabilitation, alternative behavioural therapies could be developed to focus on re-evaluating the interpretation of respiratory sensations, such as through breathing exercises (40) or mindfulness.

Our findings also demonstrate the first steps towards using FMRI as a tool for patient stratification. We speculate that this is most likely to be achieved in real life by using a detailed understanding of neural mechanisms to guide the development of appropriate behavioural tests (questionnaires, computerised tasks) that can be used at the bedside. These tests could be incorporated with appropriate clinical measures to personalise the treatment of COPD, targeting treatment options where they are needed in each individual.

### Conclusions

In conclusion, we have demonstrated that improvements in breathlessness over the course of pulmonary rehabilitation likely reflect changes in associative learning. Our findings imply that pulmonary rehabilitation shifts perceptions to become more dependent on observations and less upon learned associations, thus reducing “over-perception” of symptoms. Understanding the neural processing of breathlessness in a clinical population is crucial for advances to be made in its treatment, such as the development of patient stratification, leading to individualised treatments that may target breathlessness independently of disease mechanisms.

## SUPPLEMENTARY METHODS

### Participants

We recruited 44 (16 F, mean age 68+/-8 years) patients with COPD from the pulmonary rehabilitation services in Oxfordshire, UK. In the present study, we report findings from the 31 patients (21 male, mean age 68 +/-(SD) 9) who completed pulmonary rehabilitation.

Some of the findings have already been published (14) in a study that compared baseline (pre-rehabilitation) FMRI findings with a healthy control group. A more detailed description of the development and validation of the word-cue task has also been published(41).

Patients were studied on two separate occasions; in the two weeks prior to commencement of pulmonary rehabilitation and subsequently in the four weeks after its completion. All participants gave written, informed consent and the study was approved by Oxfordshire Research Ethics Committee A.

### MRI data acquisition

Imaging was performed at the University of Oxford Centre for Clinical Magnetic Resonance Research. Scans were always done in the same order. Participants undertook two separate FMRI scans. Each FMRI scan lasted 8 minutes and 20 seconds, with a break between scans allowing participants a chance to cough. The break was no longer than 1 minute in duration.

#### Scanner

MRI was performed with a 3 T Siemens Tim Trio scanner, with 40 mT/m gradient strength and a 12 channel receive, single channel transmit head coil (Nova Medical).

#### BOLD scanning

A T2*-weighted, gradient echo EPI was used for functional scanning. The field of view (FOV) covered the whole brain and comprised 45 slices (sequence parameters: TE, 30 ms; TR, 3 s; flip angle, 87**°**; voxel size, 3 × 3 × 3 mm; field of view,192 x192 mm; (GRAPPA factor, none) echo spacing, 0.49 ms), with 168 volumes (scan duration, 8 mins 20s) for the task fMRI.

#### Structural scanning

A T1-weighted structural scan (MPRAGE, sequence parameters: TE, 4.68 ms; TR, 2040 ms; flip angle, 8°; voxel size, 1 × 1 × 1 mm; field of view, 200 mm; inversion time, 900 ms; bandwidth; 130 Hz/Px) was acquired. This scan was used for registration of functional images.

#### Additional scanning

Fieldmap scans (sequence parameters: TE1, 5.19 ms; TE2, 7.65 ms; TR, 488.0 ms; flip angle,60°; voxel size, 3.5 × 3.5 .3.5 mm) of the B_0_ field were also acquired to assist distortion-correction.

#### FMRI task

The FMRI task comprised a set of randomised breathlessness-related cues that were presented as white text on a black background for 7 seconds. Full details of the task (and its development) can be found in previous publications (14, 41). Patients were told to rate each cue on a visual analogue scale (VAS) scale (range:0-100; anchors: ‘Not at all’ and ‘Very much’, 7 seconds), first answering the question “How breathless would this make you feel?” (wB) and second “How anxious would this make you feel?” (wA), for each given scenario. Patients were trained to reliably do the task immediately prior to the scan.

A control task to assess potential differences in baseline BOLD responsiveness between groups was employed in the form of random letter strings which were presented after every third word cue. These were not followed by any ratings and participants were instructed to ignore them. Participants were instructed to keep their eyes open for the full duration of the BOLD sequences.

#### MRI physiological measurements

Heart rate (HR) and pulse oximetry (SpO_2_, multigas monitor, model 9500, MR Equipment), respiration (respiratory bellows around the chest) and end-tidal partial pressures of oxygen (PETO_2_) and carbon dioxide (PETCO_2_; Datex, Normocap, nasal cannula (Salter Labs)) were continuously measured during the scan. All physiological data were sampled at 50Hz and recorded along with triggers for the scan volumes via PowerLab 8 (ADinstruments), using Chart5 (ADinstruments).

### Psychological measurements

#### Self report questionnaires

The following self report questionnaires were completed and were scored according to their respective manuals.

- Center for Epidemiologic Studies Depression Scale (CES-D) (15),
- State-Trait Anxiety Inventory (16)
- Fatigue Severity Scale (17),
- St George’s Hospital Respiratory Questionnaire (SGRQ) (18),
- Medical Research Council (MRC) breathlessness scale (19),
- Dyspnoea-12 (D-12) questionnaire (20),
- Catastrophic Thinking Scale in Asthma (modified by substituting the word “breathlessness” for the word “asthma”) (21)
- Pain Awareness and Vigilance Scale, (modified by substituting the word “breathlessness” for the word “pain”) (22).
- Behavioral Inhibition System/Behavioral Activation System scale(23).

The Catastrophic Thinking Scale in Asthma (21) and Pain Awareness and Vigilance Scale (22), were modified for breathlessness (see (41)), as no suitable breathlessness-specific questionnaires were available at the time of study. A later study has validated the Catastrophic Thinking Scale for COPD (42), and the resulting questionnaire is very similar, although not identical, to our questionnaire.

### Spirometry and exercise testing

Patients undertook spirometry (FEV_1_ and FVC, performed by a trained respiratory nurse) and an exercise test (Modified Shuttle Walking Test (MSWT), performed twice). HR and SpO_2_ were measured every minute (preexercise to 10 min post-exercise) using a noninvasive fingertip pulse oximeter (GO_2_, Nonin Medial Inc.). Immediately before and after the MSWT, breathlessness ratings were obtained using the modified Borg scale (43).

### Analysis of behavioural data

Questionnaires were scored by hand according to their respective manuals, with comparisons made using Student’s t-test and Spearman’s rank correlation coefficient. VAS scores were averaged and used for the FMRI analysis. For spirometry, the measurement associated with the highest FEV_1_ was used. For MSWT, the measurement associated with the largest distance was used.

**Full correlation matrices** were calculated for all behavioural and physiological measures at baseline and for the change with pulmonary rehabilitation, using MATLAB (R2013a, The Mathworks, Natick, MA). **Partial correlation matrices** were calculated on correlated variables, defined as *p* < 0.05 (uncorrected).

**Mediation analysis.** To explore the relationship between the fMRI task measures (wB and wA) the D12 questionnaire (our clinical measure of breathlessness) and other behavioural measures, we conducted a mediation analysis using the CANlab mediation toolbox (25, 44, 45).

Mediation incorporates variables that may potentially mediate a correlation (X←→Y), calculating the correlation between both variables (XM and MY). If the combination of these correlations is significant, then a variable may be deemed a potential mediator. The residuals of the regressions (XM and MY) can then be regressed to indicate whether the direct (unmediated) relationship still stands.

### FMRI Data Analysis

#### Preprocessing

Image processing was performed using the Oxford Centre for Functional Magnetic Resonance Imaging of the Brain Software Library (FMRIB, Oxford, UK; FSL version 5.0.8; http://www.fmrib.ox.ac.uk/fsl/). The following preprocessing methods were used prior to statistical analysis: motion correction and motion parameter recording (MCFLIRT(46, 47)), removal of the non-brain structures (skull and surrounding tissue) (BET(48)), spatial smoothing using a full-width half-maximum Gaussian kernel of 5 mm, and high-pass temporal filtering (Gaussian-weighted least-squares straight line fitting; 100 s). B_0_ field unwarping was conducted with a combination of FUGUE and BBR (Boundary-Based-Registration; part of FEAT: FMRI Expert Analysis Tool, version 6.0(49). Data denoising was conducted using a combination of independent components analysis (ICA) and retrospective image correction (RETROICOR)(10, 33, 50-52). Timepoints in the dataset subject to large, rapid motion artefacts were detected using FSL.

#### Image registration

After preprocessing, the functional scans were registered to the MNI152 (1×1×1mm) standard space (average T1 brain image constructed from 152 normal subjects at the Montreal Neurological Institute (MNI), Montreal, QC, Canada) using a two-step process: 1) Registration of subjects’ whole-brain EPI to T1 structural image was conducted using BBR (6 DOF) with (nonlinear) fieldmap distortion-correction (49) and 2) Registration of the subjects’ T1 structural scan to 1 mm standard space was performed using an affine transformation followed by nonlinear registration (FNIRT)(53).

#### Functional voxelwise and group analysis

Functional data processing was performed using FEAT (FMRI Expert Analysis Tool), part of FSL. The first-level analysis in FEAT incorporated a general linear model (54), with explanatory variables (EVs) at the first (individual) level for word presentation, and two EVs modelling the variability in VAS responses to the breathlessness and anxiety cues presented during scanning. Modelling this way meant that the word presentation EV was normalised, and this normalised EV was the primary measure used to examine inter-individual variability in brain responses to pulmonary rehabilitation.

Functional voxelwise analysis incorporated HRF modeling using three FLOBS regressors to account for any HRF differences caused by slice-timing delays, differences between the brainstem and cortex, or between individuals (55, 56). Time-series statistical analysis was performed using FILM, with local autocorrelation correction (57). The second and third waveforms were orthogonalised to the first to model the ‘canonical’ HRF, of which the parameter estimate was then passed up to the group analysis.

#### Higher level analyses

Group analyses were conducted using a mixed-effects analysis in FLAME 1+2 (FMRIB’s Local Analysis of Mixed Effects (58)). Z statistic images were thresholded using clusters determined by Z > 2.3 and a (corrected) cluster significance threshold of *p* < 0.05.

#### Higher level analysis 1 - Assessing variability of response to pulmonary rehabilitation

A middle level, fixed-effects analysis was conducted to calculate the difference in cope images from pre-rehab to post-rehab for each subject. These difference images were then taken up into a third, highest level, where EVs were included for mean BOLD activity, change in the VAS scales relating to “*How breathless would this make you feel?*”(wB) and “*How anxious would this make you feel?*” (wA).

#### Higher level analysis 2 – Baseline factors associated with improved breathlessness following pulmonary rehabilitation

The cope images corresponding for each subject prior to rehabilitation were taken up into a higher level analysis, where EVs were included for mean BOLD activity, pre-rehab wB, pre-rehab wA, pre-post rehab change in wB and pre-post rehab change in wA.

All FMRI data processing was carried out within FEAT (FMRI Expert Analysis Tool) Version 6.0, part of FSL (Oxford Centre for Functional Magnetic Resonance Imaging of the Brain (FMRIB) Software Library, version 5.0 www.fmrib.ox.ac.uk/fsl). The cluster Z threshold was set to 2.3 and the corrected cluster significance threshold to p=0.05 (26).

Images were registered to the MNI152 standard space using an affine registration (FMRIB Linear Image Registration Tool) between the EPI and T1 structural scan and a nonlinear registration (FMRIB NonLinear Image Registration Tool) between T1 structural scans and the MNI standard brain.

## PULMONARY REHABILITATION DETAILS

As pulmonary rehabilitation courses vary in duration and content we briefly describe the course that the patients in this study undertook. The Oxfordshire pulmonary rehabilitation programme is a cohort course which consists of two-hour sessions performed twice weekly for six weeks in an outpatient setting, that is currently (December 2016) being run by Oxford Health NHS Foundation Trust.

Outpatient settings were non-medical facilities with the appropriate exercise equipment available, e.g. sports halls. Each session consisted of 1 hour exercise (under supervision) and 1 hour education.

Exercise sessions included both aerobic and strength exercises, tailored to the individual’s ability. Aerobic exercises could include step-ups, walking (on the spot or treadmill) and exercising on a cycle ergometer. Strength exercises were conducted in sets (usually 3) of ten, and included sit-to-stand exercises, biceps curls, upright row and leg extensions.

Education sessions included items such as ‘introduction to rehabilitation’, ‘management of breathlessness’, ‘airway clearance’, ‘understanding your lung condition’, ‘home exercises’, medicine management’, ‘staying healthy’, ‘stress and relaxation’, ‘pacing and energy conservation’, ‘smoking cessation’, ‘continuing support’, ‘sexual function’ and ‘advanced care plans’. Supervised modified shuttle walking tests were used to measure patients’ improvement throughout the programme.

## ADDITIONAL RESULTS SECTION

Correlation coefficients for behavioural measures. Full and partial correlations are listed in supplementary tables 1 and 2 below. Random letter strings: No changes in brain activity to random letter strings that correlated with either wB or wA were observed.

## GROUP MEAN WORD-CUE ACTIVITY

**Supplementary Figure.**
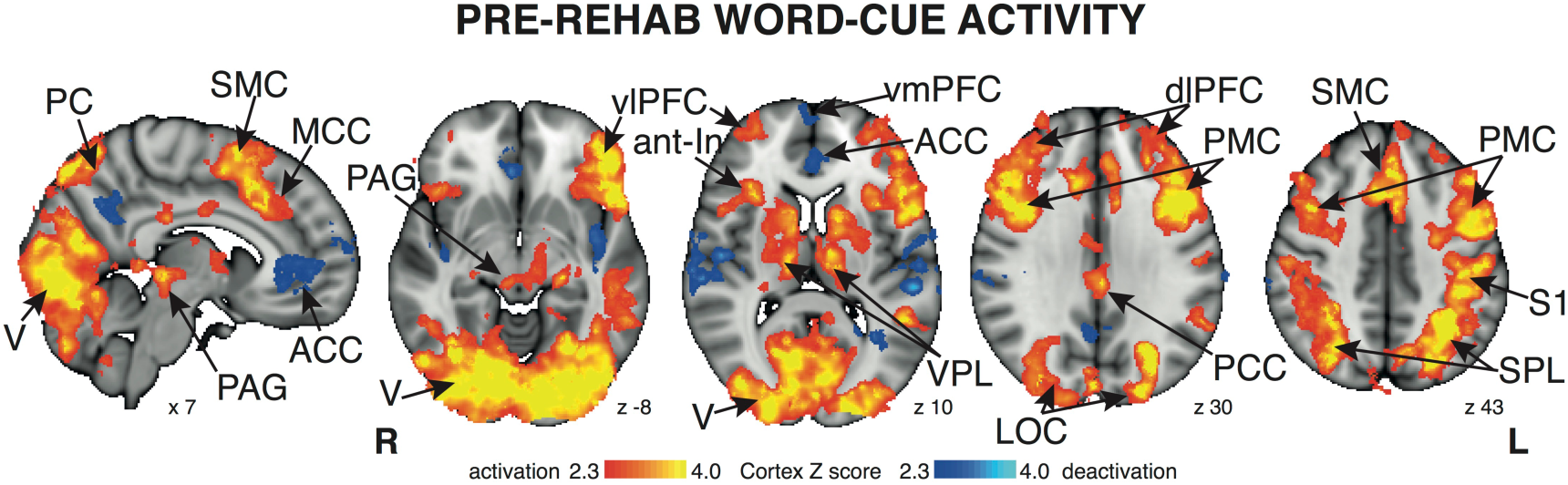
Mean BOLD response to presentation of word cues prior to pulmonary rehabilitation (not weighted by individual measures of wB and wA). The images consist of a colour-rendered statistical map superimposed on a standard (MNI 1×1×1 mm) brain, and significant regions are displayed with a threshold Z > 2.3, with a cluster probability threshold of p < 0.05 (corrected for multiple comparisons). Abbreviations: vlPFC, ventrolateral prefrontal cortex; vmPFC, ventromedial prefrontal cortex; ACC, anterior cingulate cortex; MCC, middle cingulate cortex; PCC, posterior cingulate cortex; SMC, supplementary motor cortex; PMC, premotor cortex; PC, precuneus cortex; ant-In, anterior insula; PAG, periaqueductal gray; V, visual cortex; LOC, lateral occipital cortex; VPL, ventral posterolateral thalamic nucleus; S1, primary sensory cortex; SPL, superior parietal lobule; L, left; R, right; activation, increase in BOLD signal; deactivation, decrease in BOLD signal.

**Supplementary Figure:**
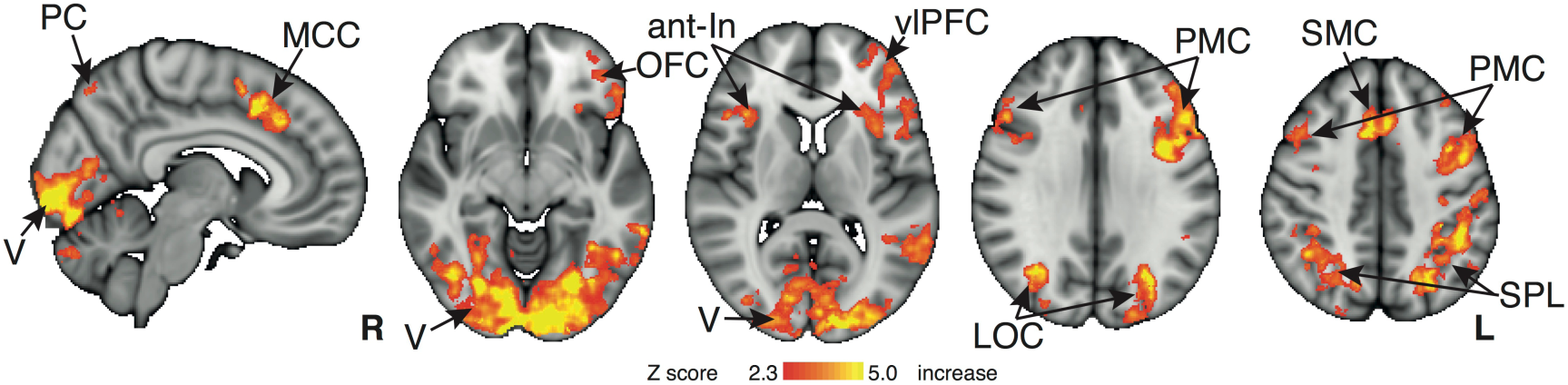
Group mean BOLD response to presentation of word cues following pulmonary rehabilitation (not weighted by individual measures of wB and wA). The images consist of a colour-rendered statistical map superimposed on a standard (MNI 1×1×1 mm) brain, and significant regions are displayed with a threshold Z > 2.3, with a cluster probability threshold of p < 0.05 (corrected for multiple comparisons). Abbreviations: vlPFC, ventrolateral prefrontal cortex; OFC, orbitofrontal cortex; MCC, middle cingulate cortex; SMC, supplementary motor cortex; PMC, premotor cortex; PC, precuneus cortex; ant-In, anterior insula; V, visual cortex; LOC, lateral occipital cortex; SPL, superior parietal lobule; L, left; R, right.

### Full Correlation Matrix for behavioural scores prior to pulmonary rehabilitation

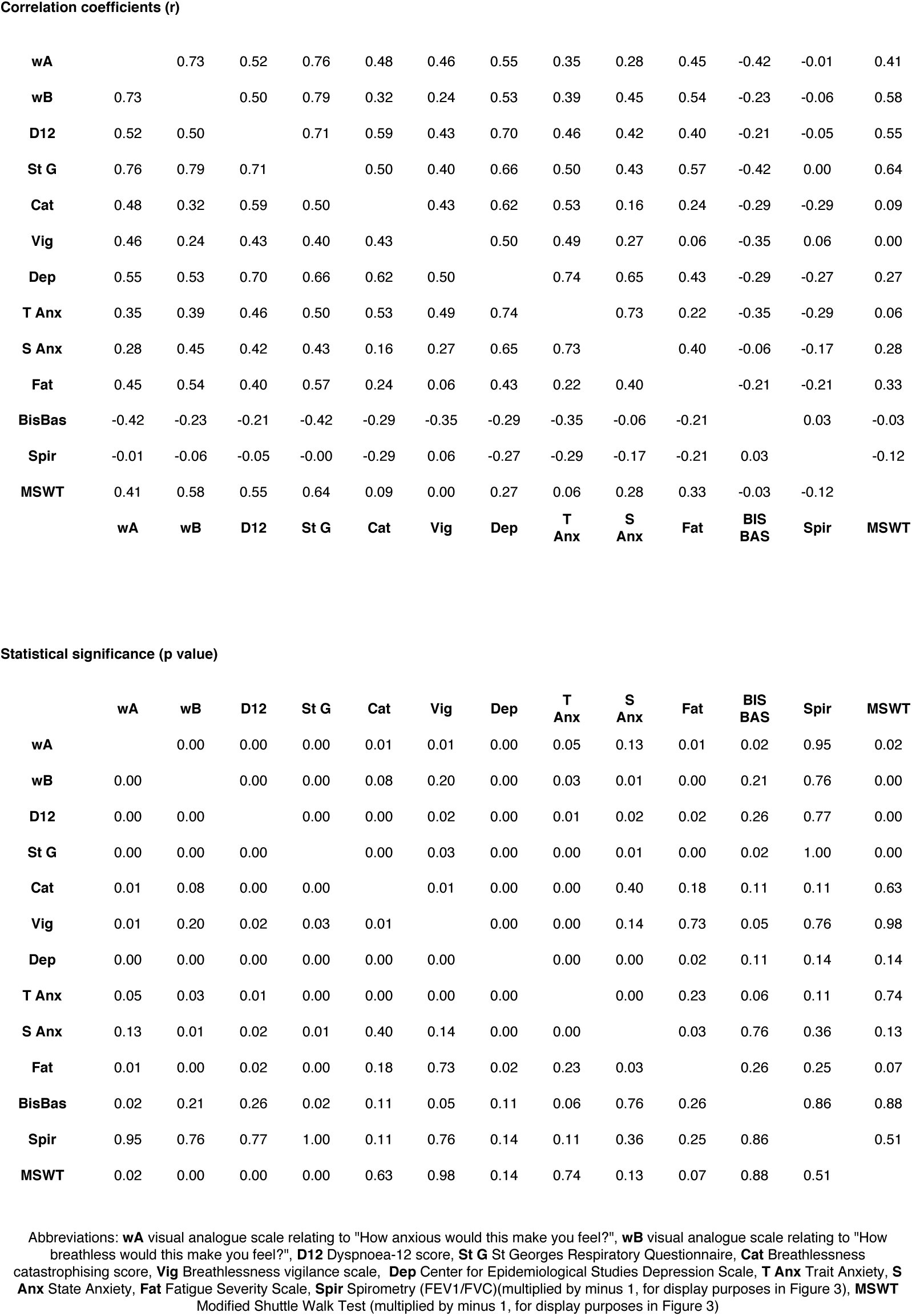
Full Correlation Matrix for behavioural scores prior to pulmonary rehabilitation

### Partial correlation matrix for behavioural scores prior to pulmonary rehabilitation

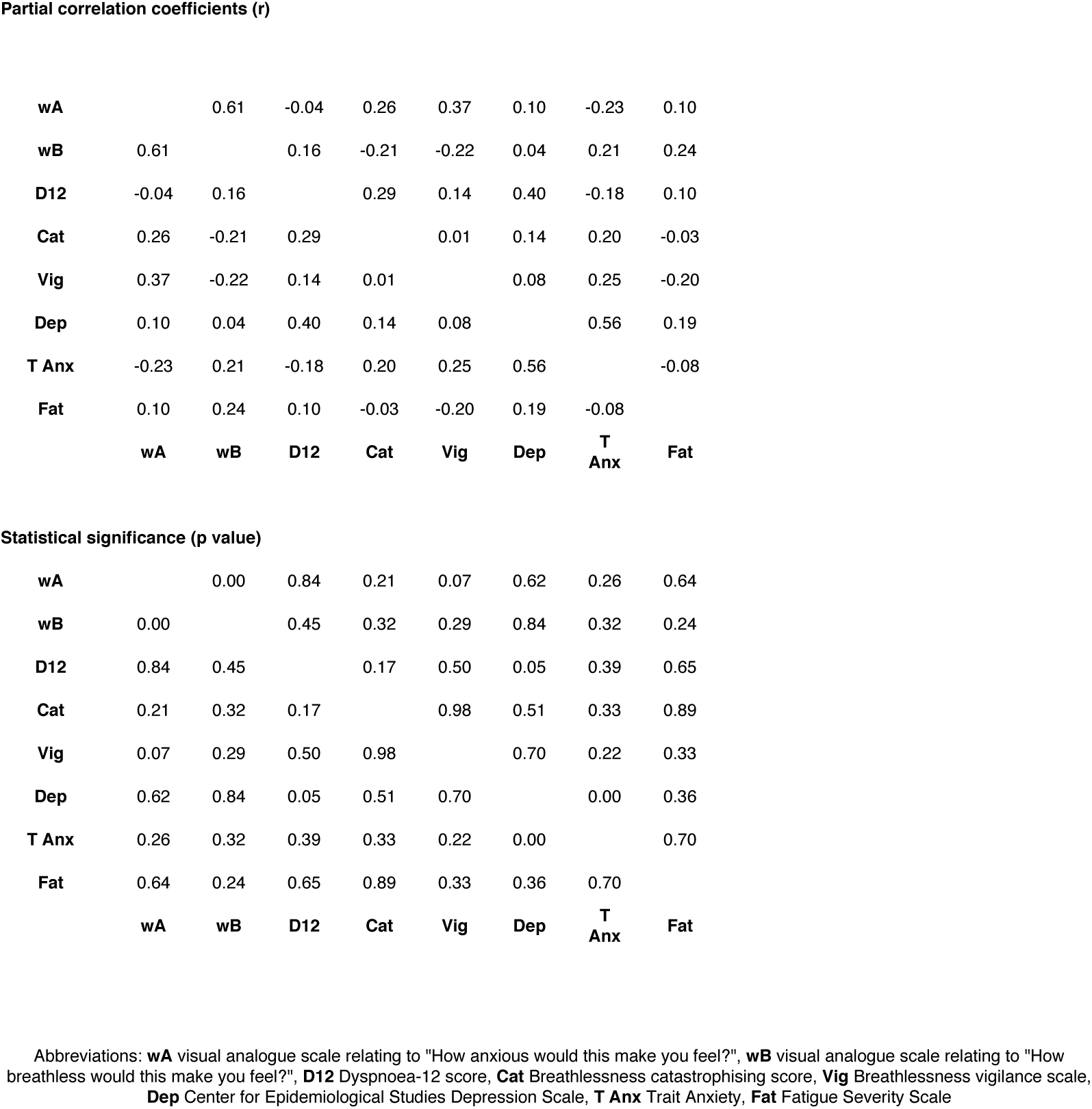
Partial correlation matrix for behavioural scores prior to pulmonary rehabilitation

### Full Correlation Matrix for pre vs post rehabilitation scores

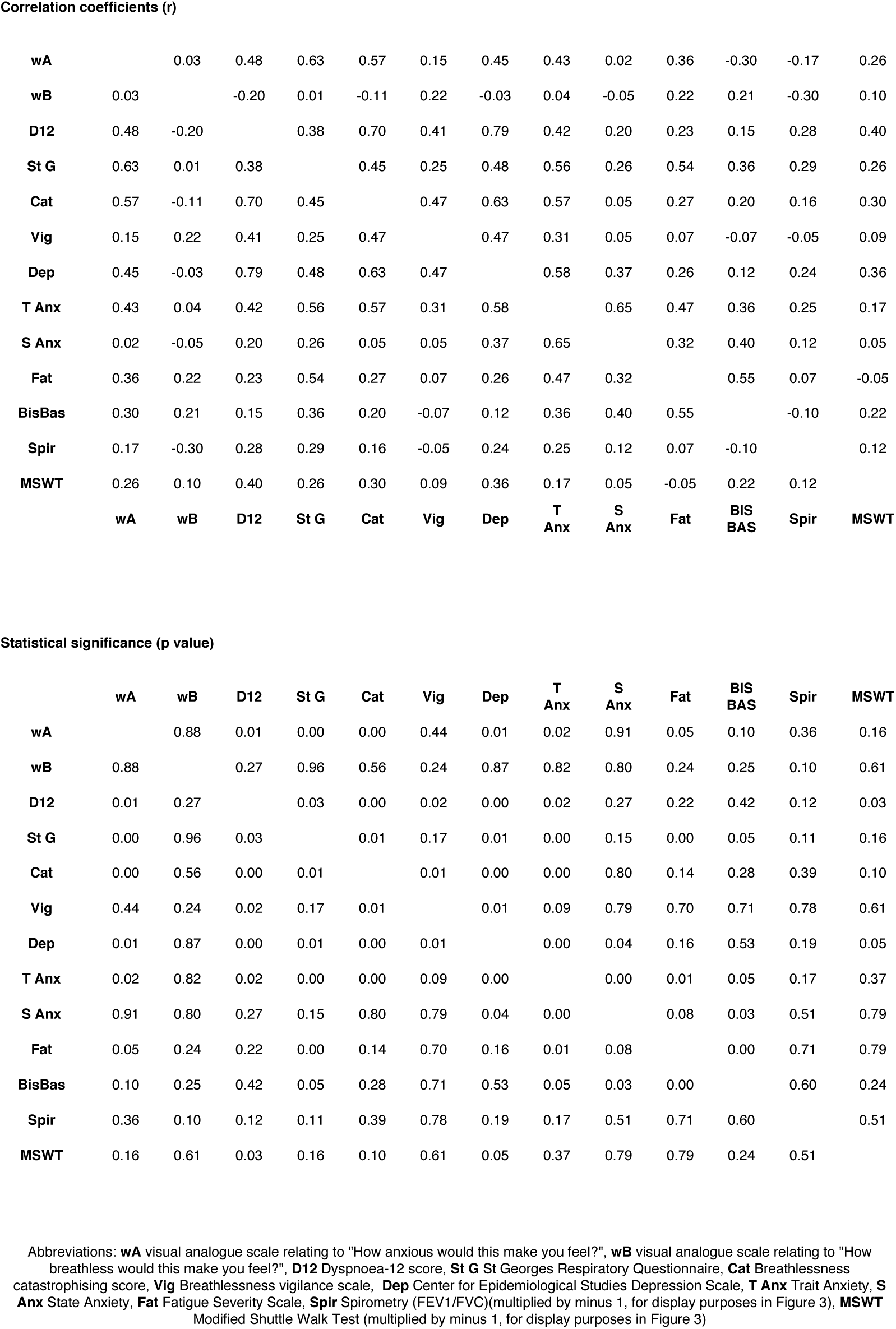
Full Correlation Matrix for pre vs post rehabilitation scores

### Partial Correlation Matrix for pre vs post rehabilitation scores

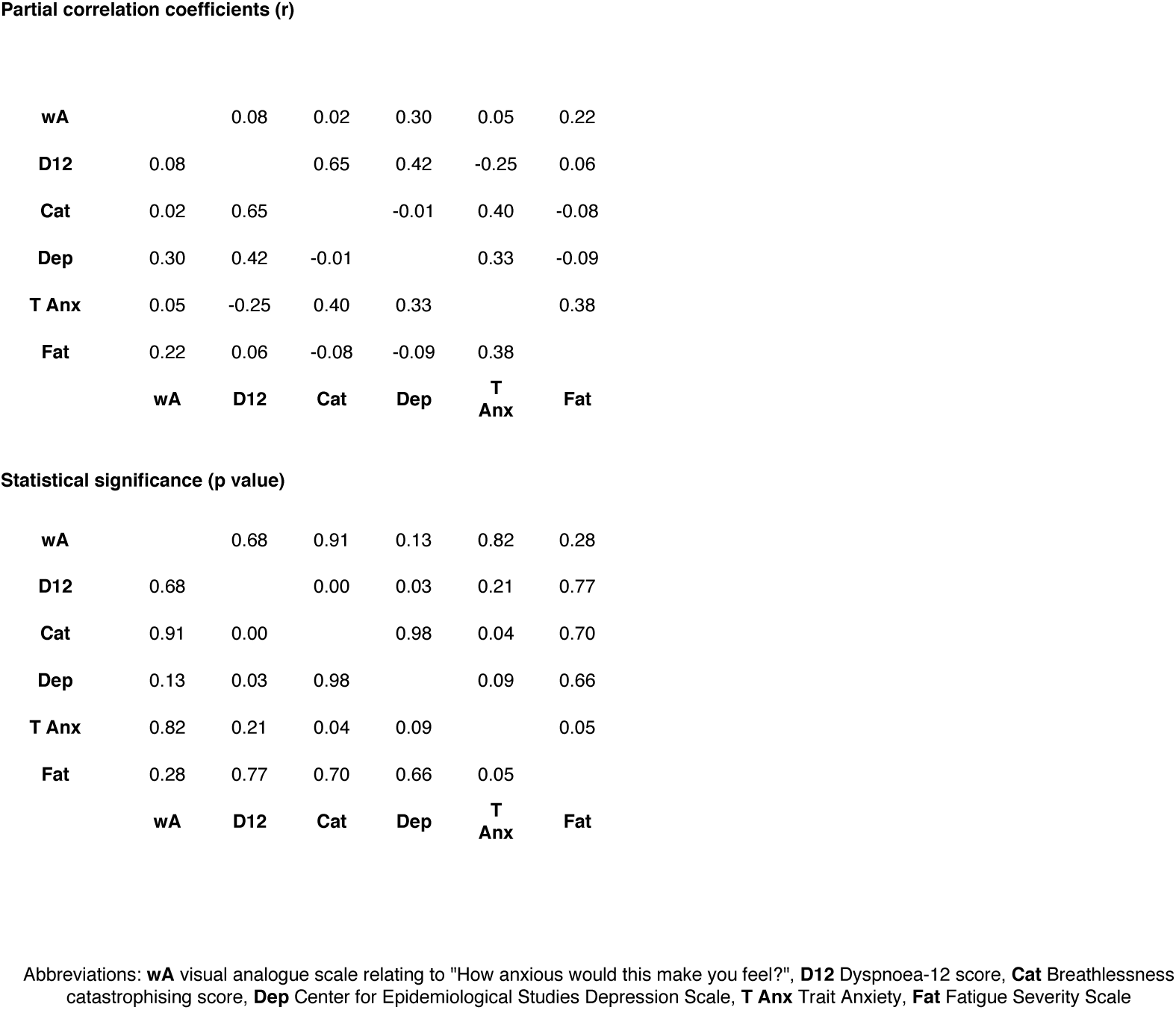
Partial Correlation Matrix for pre vs post rehabilitation scores

## ACKNOWLEDGEMENTS

The authors thank Steve Knight, Bethan Hughes, Isabel Chabata, Debby Nicoll, Tara Harris, Fran Sinfield, Emma Tucker, Rachel Lardner, Julie Young and the Oxfordshire Pulmonary Rehabilitation Team for their continued support of the study; and Richard Wise, Andrea Reinecke, Rob Davies, Annabel Nickol and Najib Rahman for their generous input. We wish to thank Caroline Jolley, Sara Abdallah, Sara Booth and Ben Ainsworth for their comments on previous versions of this manuscript. This study was supported by the Medical Research Council (UK) and the National Institute for Health Research (NIHR) Oxford Biomedical Research Centre based at The Oxford University Hospitals Trust, Oxford.

